# Optogenetic silencing of immature and mature neurons in dentate gyrus to assess their roles in memory discriminations

**DOI:** 10.1101/335992

**Authors:** Milenna T. van Dijk, Zejia Angel Yu, Younghun Lim, René Hen, André A. Fenton

## Abstract

Discriminating similar memories and events depends on the dentate gyrus region of the hippocampus. This region is also distinctive because neurogenesis continues in adulthood. Whether both mature and immature granule cells play a role in memory discrimination, and whether the roles are distinct is actively investigated. Here we demonstrate that manipulating either mature or immature granule cells can impair discrimination of similar active place avoidance memories, but the manipulations have different effects. We also observe that prior experience modulates which memories are compromised by inactivation of immature neurons. These data demonstrate the importance of the dentate gyrus network of cells for memory discrimination.

## INTRODUCTION

To accurately encode and store new memories, it is crucial to discriminate between new unique events and familiar repeating experiences that should be treated as memories to update. Pattern separation is the neural computation that makes two or more outputs of a system more distinctive than the corresponding input patterns. Pattern separation is proposed to underlie the ability to discriminate memories of similar events. Computational theory (Marr, 1971; Treves and Rolls, 1994) and anatomy (Amaral et al., 1990; Amaral et al., 2007; Mulders et al., 1997) pointed to the dentate gyrus (DG) as a site for pattern separation within the hippocampus circuit. Because the DG has 4-5 times more principal neurons than its upstream and downstream regions, it is structurally poised to sparsify and orthogonalize the information that is carried by the divergent afferents from entorhinal cortex. DG is also functionally poised for pattern separation, as granule cells, the DG’s principal excitatory neurons, have low, less than 1%, interconnectivity (Amaral et al., 2007) and they discharge infrequently due to strong inhibition and hyperpolarized resting membrane potential (Coulter and Carlson, 2007; Mody, 2005). Consistent with a role in memory discrimination, manipulations that compromise DG function tend to impair difficult memory discriminations (Burghardt et al., 2012; Gilbert et al., 2001; Kheirbek et al., 2013; Lee et al., 2005; McHugh et al., 2007; Nakashiba et al., 2012).

The DG is one of a few brain regions in which, during adulthood, neurons are born, proliferate, and integrate into the network functionally. Immature neurons incorporate into the hippocampal network two weeks after mitosis (Gu et al., 2012; Toni et al., 2008; Zhao et al., 2006), but these cells remain distinct from mature neurons in their electrophysiological function for at least an additional six weeks (Overstreet-Wadiche and Westbrook, 2006; Zhao et al., 2006). Immature granule cells are more excitable than mature granule cells, in part due to reduced feedback inhibition (Danielson et al., 2016; Marin-Burgin et al., 2012) and they show enhanced synaptic plasticity (Ge et al., 2007; Schmidt-Hieber et al., 2004). During active behaviors, immature neurons are more easily excited, discharging in less discriminating spatial fields, and their activation is more similar across different situations than the discharge of mature granule cells (Danielson et al., 2016; Marin-Burgin et al., 2012). Nonetheless, studies indicate that adult neurogenesis significantly contributes to DG-dependent memory discrimination and the ability to resolve interference between potentially conflicting experiences (Arruda-Carvalho et al., 2011; Burghardt et al., 2012; Clelland et al., 2009; Danielson et al., 2016; Kheirbek et al., 2012; Sahay et al., 2011a).

Using an active place avoidance task, we previously demonstrated that immature granule cells are crucial for memory discrimination and cognitive flexibility (Burghardt et al., 2012). That study and other findings suggest an indirect role for immature granule cells in memory by regulating the inhibition of mature granule cells (Anacker et al., 2018; Burghardt et al., 2012; Drew et al., 2016; Lacefield et al., 2010; Park et al., 2015; Sahay et al., 2011a), whereas other studies have instead suggested that immature neurons are directly involved in memory encoding (Aimone et al., 2010b; Aimone et al., 2009; Rangel et al., 2013).

The Burghardt et al. (2012) study chronically ablated adult neurogenesis and although it was comprehensive, during the post-lesion months the hippocampus could have reorganized to adjust to the circumstances of the new computational constraint. This and other difficulties of interpreting the results of permanent lesions compromises our ability to infer the role of immature neurons (Bures and Buresova, 1990). Additionally, as with any complex process, the active place avoidance task likely requires multistage processing and immature neurons might only contribute to specific stages, for example encoding but not retrieval, or *vice versa*. Permanent ablation of adult neurogenesis cannot discriminate between a selective and a general role in a multistage process, therefore to determine if immature neurons have a selective role in memory discrimination, we exploit the cell-specific reversible lesion approach that is offered by optogenetics. The present investigations used two mouse strains that express Cre recombinase. We bred each strain with a mouse expressing a floxed stop codon that controls expression of an inhibitory opsin. POMC-Cre mice express Cre mostly in dentate gyrus granule cells. Although Cre is also expressed in additional regions, we get functional specificity of the inactivation by targeting the light to the DG. Nestin-CreERT2 mice express Cre in immature granule cells selectively after tamoxifen administration. We then used DG-targeted laser inhibition of immature neurons to selectively inhibit immature granule cells during encoding and/or retrieval of discriminative active place avoidance memories.

## METHODS

### Animals

Sixteen POMC-Cre-Halorhodopsin male adult mice (8 Cre^+^ and 8 Cre^−^) and 44 Cre-Nestin-ERT2-Halorhodopsin adult mice were used (males: 15 Cre^+^ and 13 Cre^−^; females: 7 Cre^+^ and 9 Cre^−^). Twelve NestinCreERT2-Arch male adult mice were used for the object tasks (6 Cre^+^ and 6 Cre^−^). All procedures and care adhered to protocols approved by New York University Animal Welfare Committee, which follow NIH guidelines. POMC-Cre-Halorhodopsin mice result from the following cross: B6.FVB-Tg(Pomc-cre)1Lowl/J (RRID:IMSR_JAX:010714) x B6;129S-Gt(ROSA)26Sor^tm39(CAG-hop/EYFP)Hze^/J (RRID:IMSR_JAX:014539) and NestinCreERT2-Halorhodopsin mice result from crossing NestinCreERT2 (Dranovsky et al., 2011) x B6;129S-Gt(ROSA)26Sor^tm39(CAG-hop/EYFP)Hze^/J (for Halorhodopsin; RRID:IMSR_JAX:014539) or B6;129S-Gt(ROSA)26Sor^tm35.1(CAG-aop3/GFP)Hze^/J (for Archaerhodopsin; RRID:IMSR_JAX:012735). At 8 weeks NestinCreERT2-Halorhodopsin mice and littermate controls and NestinCreERT2-Arch and littermate controls were injected with 125μl tamoxifen daily for 5 days. Two-three weeks later mice underwent surgery for bilateral optic fiber placement and five weeks post injection mice started the behavioral paradigm.

### Surgery

Mice were anesthetized with isoflurane (3% for induction 1.5-1.75% maintenance) for surgical procedures. Ketoprofen analgesia was administered for three days after surgery. A minimum of two weeks passed before experimental procedures began. Bilateral optic fibers (0.125 diameter optic fibers (NA 0.37) in 1.25mm zirconia ferrules) were surgically implanted at ML: ±1.25, AP: −1.8, DV −1.55 using C&B Metabond to attach the ferrules to the skull.

### Active place avoidance Pretraining-Initial-Conflict (PIC) protocol

A commercial active place avoidance apparatus and software was used (Bio-Signal Group, Corp., Acton, MA). Mice were trained on the active place avoidance task in a black-curtained room with distinctive visual cues on the curtains. For optogenetic silencing, a 593 nm laser was attached to the implanted optic fibers. When the laser was on, the peak power in the brain was 15 mW. On day 1, mice explored the rotating disk (0.75 rotation/min) during a 30-minute Pretraining trial with no shock. On day 2 mice learned to avoid the Initial location of a computer-controlled 0.2 mA, 60 Hz, 500 ms foot shock in a 60° unmarked sector-shaped zone during two 30-minute training trials (2 hours apart). The shock zone was stationary with respect to the room (Cimadevilla et al., 2001). On the third trial of day 2 the mice were trained in a Conflict trial with the shock zone relocated 180°. Mice were euthanized and perfused 105 minutes after the start of the conflict trial.

Eight male transgenic POMCCre^+^-Halorhodopsin mice that express the light sensitive chloride pump halorhodopsin selectively in DG granule cells and eight male Cre^−^ littermate controls received the active place avoidance training as described above with the laser on for every trial.

Eight Nestin-CreERT2^+^-Halorhodopsin and seven Cre^−^ littermate controls received the same PIC training protocol with the laser on for every trial. Thirteen NestinCreERT2^+^-Halorhodopsin and fifteen NestinCreERT2^−^ littermate controls received the PIC training protocol with the laser only on for the Conflict trial.

### Active place avoidance two-zone task

A week after the start of the PIC protocol mice received 2 days of additional training in a two-zone active place avoidance protocol. In this protocol a rotating arena frame shock zone is added to the already learned stationary room frame shock zone. To avoid shock mice have to learn to escape through the center where there is no shock and go to the other side. This is a difficult task for all mice to learn. Therefore there were three 30 min trials a day for two days. Most mice had learned to avoid the shock zones by the final trial.

### Object tasks

Six male adult NestinCreERT2^+^-Arch and six littermate controls underwent two days of training in the object tasks. The environment was the same circular arena used for training in the PIC protocol, but the enclosure was a square box with cues on three out of four walls surrounding it and a plastic floor.

#### Novel object task

On day 1 mice had a ten minute habituation session of the environment with no objects. After the habituation trial, the mice had 2 hours of home cage rest. Two equivalent objects were then placed equidistant from the center and the walls of the circular enclosure. Subsequently, the mice spent 10 minutes in the enclosure to explore the objects. After another two hours of rest, the object that the mouse had spent the least time exploring was replaced by a novel object, similar in height but a different shape and material. Between mice, objects and arena were sprayed with alcohol to get rid of odors.

#### Object pattern separation task

On day 2 mice again did a 10 min habituation trial in the same environment. After two hours of rest, mice explored two equivalent objects, placed equidistant from the center and walls symmetrically for 10 minutes. Finally, after another two hours of rest, the object the mice had explored the least was displaced 4 cm away from the center.

#### Analysis of object tasks

Videos were recorded of each mouse’s’ behavior. Analysis was offline. The time that the animal explored each object was measured twice per trial per animal and averaged. To construct difference scores, the time spent with the familiar object was subtracted from time spent with the novel or displaced object and this difference was normalized by the total time spent exploring both objects.

### Histology

Mice were anesthetized with sodium pentobarbital (100mg/kg, i.p.) and perfused with saline followed by 4% paraformaldehyde in 0.1 M phosphate buffer. Brains were postfixed overnight. The next day, brains were cryoprotected in 30% sucrose 0.1 M phosphate buffer solution. Sections (35 μm) were incubated in c-Fos Anti-Rabbit primary antibody (1:8000; Millipore: RRID:AB_2631318) and secondary 594 nm Goat Anti-Rabbit Alexa Fluor (1:500; ThermoFisher Scientific: RRID:AB_2556545) for fluorescent microscopy. Sections were mounted using Vectashield mounting medium with DAPI (Vector Labs) and coverslipped. Image acquisition occurred with an Olympus VS120 Virtual Slide Microscope with a 20x 0.75 NA objective. Composite images of DAPI and c-Fos immunofluorescence (excitation wavelengths: 455 nm and 580 nm) are displayed. Neuronal activity during the conflict trial was estimated by counting c-Fos+ cells in six slices per animal.

## RESULTS

### Selective impairments after optogenetic silencing of DG granule cells

To test whether acutely silencing granule cells impairs spatial memory discrimination, we had mice perform the Pretraining-Initial-Conflict avoidance learning protocol of the active place avoidance task. The PIC protocol is shown in Fig. 1A and the laser targeting the DG was on for all trials to mimic permanent functional ablation of DG neurons with limited opportunity for circuit reorganization. The mice are placed on the rotating circular arena for 30 min per trial. During the *Pretraining* session on day 1, there is no shock allowing the mouse to acclimatize to the experimental conditions. The next day, during two *Initial* training trials, mice learn to avoid a shock zone that is stationary with respect to the room. On the third *Conflict* trial, the shock zone is relocated 180° and the mice are challenged to discriminate between the initial and current (conflict) locations of shock. Note that the physical environment is identical across all trials except during the 500 ms that the mouse might experience shock, which in the aggregate is typically less than 1% of the trial (1% corresponds to 36 shocks in 30 min).

**Fig. 1.**
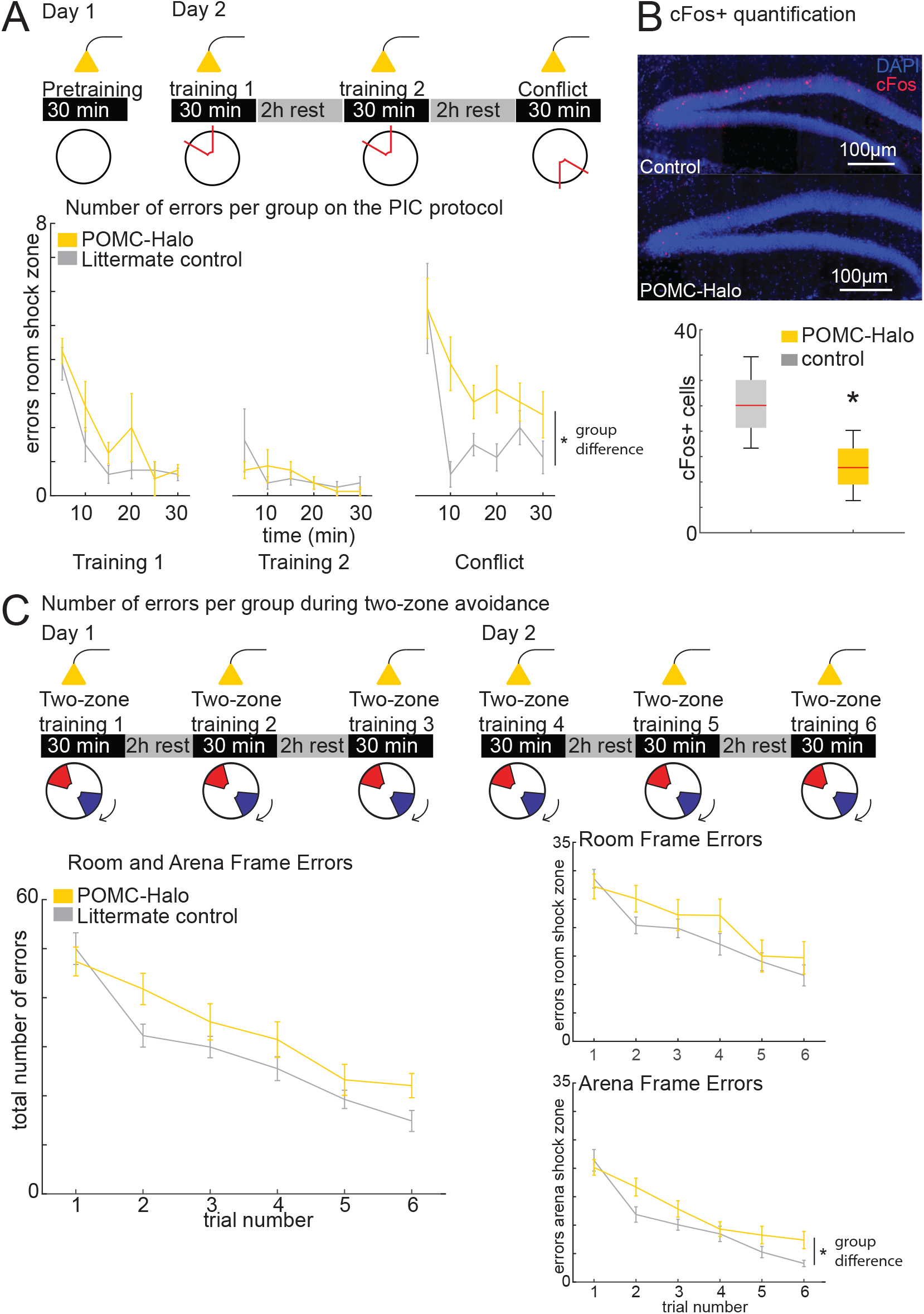
Disrupting the DG by optogenetically silencing DG granule cells impairs memory discrimination but not learning or memory. A) Top: PIC experimental protocol. Bottom: POMC-Cre^+^ mice with optogenetically-silenced granule cells are only impaired on the Conflict trial, demonstrating impaired memory discrimination. (Initial training 1: Genotype x Time repeated measures ANOVA; genotype: F_1,14_ = 2.20, p = 0.16; time: F_5,70_ = 25.0, p < 0.0001; interaction: F_5,70_ = 0.76, p = 0.58. Initial training 2: Genotype x Time repeated measures ANOVA; genotype: F_1,14_ = 0.23, p = 0.64; time: F_5,70_ = 2.14, p = 0.07; interaction: F_5,70_ = 0.89, p = 0.49. Conflict trial: Genotype x Time repeated measures ANOVA; genotype: F_1,14_ = 7.16, p = 0.02; time: F_5,70_ = 10.0, p < 0.0001; interaction: F_5,70_ = 1.61, p = 0.17). B) The number of c-Fos positive cells per section that were activated during the Conflict trial is significantly reduced in the light-silenced mice compared to the POMC-Cre^−^ control mice. C) During the two-zone task variant, mice with optogenetically silenced granule cells are impaired relative to control mice, particularly in avoiding the rotating arena shock zone. There were no genotype differences on the combined number of room and arena frame errors (Genotype X Trial repeated measures ANOVA; genotype: F_1,12_ = 0.43, p = 0.53; trial: F_5,8_ = 10.6, p=0.002; interaction: F_5,8_ = 0.24, p = 0.94), nor for stationary room frame only errors (Genotype X Trial repeated measures ANOVA; genotype: F_1,12_ = 0.02, p = 0.88; trial: F_5,8_ =3.37, p=0.06; interaction: F_5,8_ = 0.34, p = 0.87). However, there were genotype differences in the rotating arena frame errors (Genotype X Trial repeated measures ANOVA; genotype: F_1,12_ = 6.91, p = 0.02; trial: F_5,8_ =13.46, p=0.001; interaction: F_5,8_ = 0.12, p = 0.98).

Optogenetically silencing granule cells in POMC-Cre^+^ mice is demonstrated to perturb other DG cell types such as mossy cells and inhibitory interneurons through secondary, network effects (Senzai and Buzsáki, 2017). Nonetheless, the inhibition did not impair learning or memory measured on the Initial training trials, compared to POMC-Cre^−^ mice (Fig. 1A). However, when the shock zone was relocated to test memory discrimination on the Conflict trial, the light-inhibition impaired the POMC-Cre^+^ mice compared to the POMC-Cre^−^ mice (Fig. 1A). The two-way Genotype x Protocol phase, repeated measures ANOVA comparing the genotypes across the Initial training 1, Initial training 2 and Conflict training trials confirmed a selective impairment on the Conflict trial (genotype: F_1,14_ = 11.7, p = 0.004; protocol phase: F_1.42,19.9_ = 46.4, p<0.0001; interaction: F_1.42,19.9_ = 4.9, p = 0.03). Post-hoc tests show POMC-Cre^+^ mice make more errors than POMC-Cre^−^ only on the Conflict trial. After adding the shock zone to initiate Initial training 1, mice have to make a qualitative memory discrimination between the absence of shock in the prior Pretraining trial and the experience of shock during the training trials. According to our results, this memory discrimination can be accomplished after disrupting DG function. In contrast, when the shock is relocated 180° for Conflict training, the mice have to perform a quantitative memory discrimination between memories for the two locations of shock, everything else being identical. Immediate early gene analysis demonstrated that POMC-Cre^+^ mice had fewer c-Fos^+^ granule cells labeled during the conflict trial (t_6_ = 2.7, p = 0.04; Fig.1B), confirming a significant, albeit incomplete inhibition. Thus, optogenetic silencing targeting granule cells is sufficient to disrupt memory discrimination but not memory itself in the active place avoidance paradigm. A week later, the same mice were trained in a different active place avoidance task variant, in which a rotating arena-defined shock zone was added to the stationary shock zone. In this two-zone task, the mice have to run through the center of the arena to the other side as the rotating and stationary shock zones converge on every complete rotation. The mice must now make within-trial memory discriminations between the stationary and rotating locations of shock. After learning to avoid a stationary shock zone in the PIC protocol, this task can be conceptualized as a negative priming challenge because the mice had never experienced a particular arena location as unsafe, and now a specific arena zone is consistently punished by shock. This is a demanding task with substantially increased demand for cognitive control (Kelemen and Fenton, 2010), and when specifically presented in a negative priming sequence, the two-zone task variant is impaired by X-ray ablation of adult neurogenesis (Burghardt et al., 2012).

Mice were trained in the two-zone task variant during six 30-minute trials spread out over two days (Fig 1C). Note that this protocol requires the mice to use an explicit form of within-session pattern separation; mice discriminate between room-defined locations and arena-defined locations and hippocampus place cell ensemble recordings show that the cells collectively alternate every few seconds between stationary and rotating discharge patterns to represent stationary and rotating locations, respectively (Kelemen and Fenton, 2010; Kelemen and Fenton, 2016; Talbot et al., 2018; van Dijk and Fenton, 2018). The combined number of stationary and rotating shock zone errors decreased with training and the number of errors in the stationary room frame zone was similar in the two genotypes across all trials. In contrast, the optogenetic silencing of granule cells caused mice to more often enter the rotating arena frame zone, committing more errors throughout the two-zone training.

### Selective impairments after optogenetically silencing immature DG granule cells

Next, we tested the impact of selectively silencing immature granule cells (<5 weeks post-mitosis) on active place avoidance learning, memory and cognitive flexibility. The PIC protocol was the same as above, and we tested both male and female NestinCreERT2^+^-Halorhodopsin mice that conditionally express halorhodopsin in immature neurons and NestinCreERT2^−^ littermate control mice. The mice were tested five weeks after conditional recombination was induced by five days of tamoxifen administration. No differences in the number of errors were found for any of the trials between male and female mice (sex X protocol phase repeated measures ANOVA: sex: F_1,14_ = 0.83, p = 0.37; protocol phase: F_2,13_ = 55.3, p <0.0001; interaction: F_2,13_ = 0.005, p = 0.996). We collapsed the data over sex in the following analyses.

During the first and second Initial training sessions, there were no differences in errors between the genotypes (Fig. 2A). Mice of both genotypes learned and remembered and made fewer errors in the second training trial than in the first training trial (Fig. 2A). We then tested mice on the Conflict trial, which requires mice to discriminate between the previously learned shock zone location and the new location 180° away. There were no genotype differences in performance on the Conflict task, suggesting that immature neurons are not necessary for this memory discrimination. The lack of genotype differences was confirmed with a Genotype X Protocol phase (Initial training 1, Initial training 2, Conflict) repeated measures ANOVA (genotype: F_1,14_=0.16, p = 0.70; protocol phase: F_1.28,17.9_ = 88.25, p<0.0001; interaction: F_1.28,17.9_ = 2.52, p = 0.12).

**Fig. 2.**
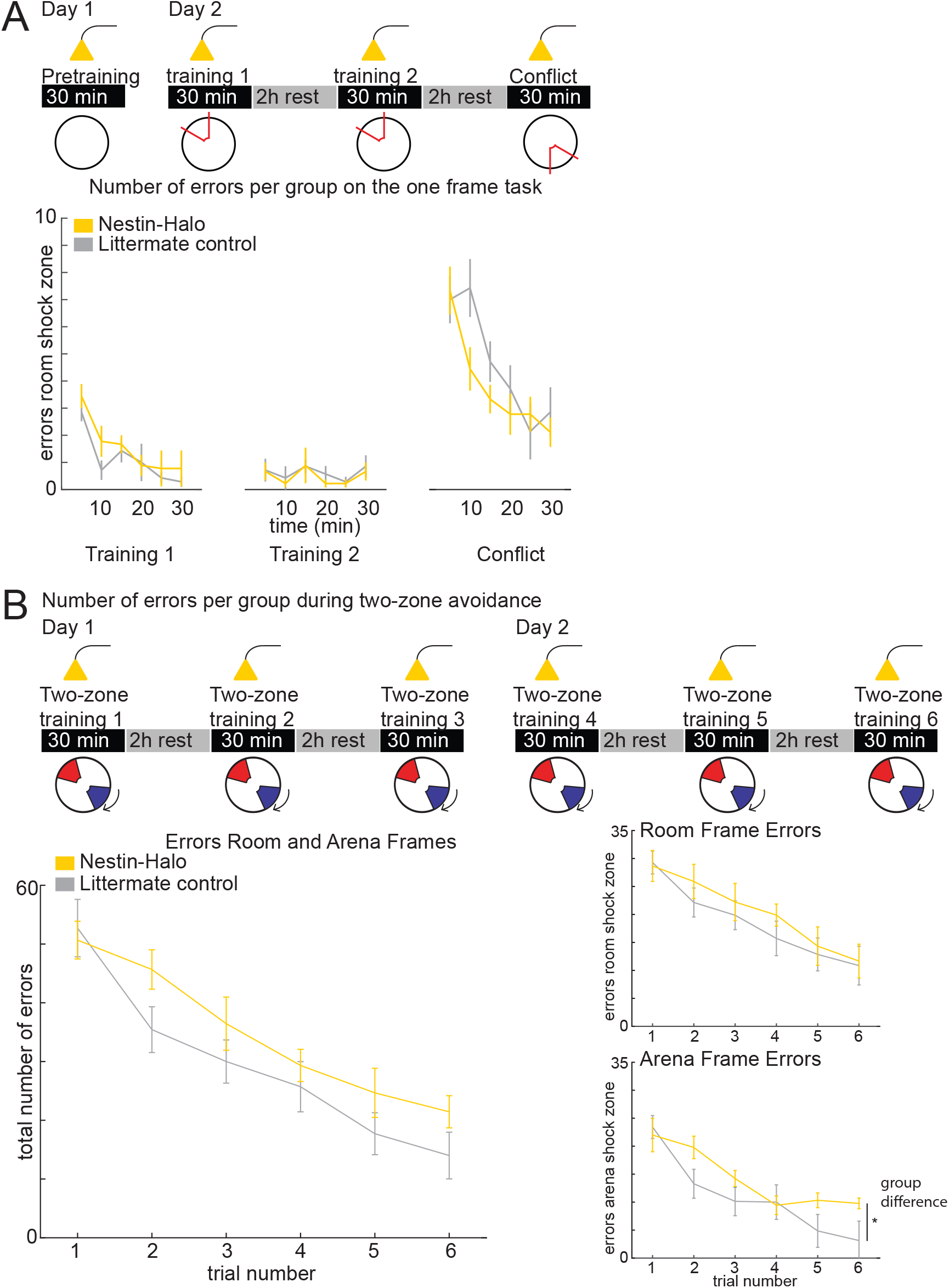
Mice with light-silenced immature granule cells are impaired on a memory discrimination task with large cognitive load, specifically to avoid the rotating shock zone. A) On a protocol with the laser on during all trials, the NestinCre^+^ mice are not impaired on the PIC protocol relative to the NestinCre^−^ mice. Initial training 1: Genotype X Time repeated measures ANOVA; genotype: F_1,14_ = 0.78, p = 0.39; time: F_5,70_ = 13.0, p < 0.0001; interaction: F_5,70_ = 0.53, p = 0.75. Initial training 2: Genotype x Time repeated measures ANOVA; genotype: F_1,14_ = 0.23, p = 0.64; time: F_5,70_ = 1.35, p = 0.25; interaction: F_5,70_ = 0.10, p = 0.99. Conflict trial: Genotype x Time repeated measures ANOVA; genotype: F_1,14_ = 1.20, p = 0.29; time: F_5,70_ = 17.87, p < 0.0001; interaction: F_5,70_ = 2.08, p = 0.09. B) One week later, in the two-zone task variant with larger cognitive load, mice with silenced immature granule cells demonstrate decreased ability to discriminate between a previously learned stationary shock zone and a new qualitatively different, rotating shock zone. There were no genotype differences on the combined number of room and arena frame errors (Genotype X Trial repeated measures ANOVA; genotype: F_1,14_ = 2.01, p = 0.18; trial: F_5,10_ = 30.3, p < 0.0001; interaction: F_5,10_ = 0.77, p = 0.59), nor for stationary room frame errors (Genotype X Trial repeated measures ANOVA; genotype: F_1,14_ = 0.39, p = 0.54; trial: F_5,10_ = 7.52, p = 0.004; interaction: F_5,10_ = 0.57, p = 0.73). However, there were genotype differences in the rotating arena frame errors (Genotype X Trial repeated measures ANOVA genotype: F_1,14_ = 4.84, p = 0.045; trial: F_5,10_ = 34.7, p < 0.0001; interaction: F_5,10_ = 3.98, p = 0.03).

It seems that optogenetic silencing of immature neurons may not be sufficient to impair DG-dependent memory discrimination, even though an impairment was previously reported for permanent ablation of neurogenesis (Burghardt et al., 2012). Furthermore, contextual fear discrimination in NestinCreERT^+^ mice was selectively impaired when immature neurons were optogenetically-silenced during encoding of an ambiguous safe chamber, but not when immature neurons were silenced during acquisition of fear conditioning in the conditioning chamber that had already been encoded (Danielson et al., 2016). It is then possible that the present optogenetic silencing either affected an insufficient number of immature neurons, or alternatively, immature neurons may play a significant role only in the DG-dependent memory discrimination trial, but this was obscured by light-inhibition of these cells during the training trials. If the cells are not active during encoding they are unlikely to be part of the representation of the initial location of the shock; consequently, they may not have been able to play a significant role in discriminating between the initial and relocated locations of the shock.

Therefore, a second cohort of NestinCreERT^+^-Halorhodopsin and NestinCreERT^−^ littermate controls were tested on the PIC protocol, this time with the laser on <u>only</u> for the DG-dependent Conflict trial. Again, there were no differences between sexes (sex X protocol phase, repeated measures ANOVA: sex: F_1,26_ = 0.32, p = 0.58; protocol phase: F_2,25_ = 28.8, p <0.0001; interaction: F_2,25_ = 0.20, p = 0.82). We collapsed the data over sex for further analysis.

Both genotypes learned to avoid and remember the stationary shock zone in the Initial training trials, and there were again no differences between genotypes on the Conflict trial (Genotype X Protocol phase repeated measures ANOVA on errors: genotype: F1,26 = 0.31, p = 0.58; protocol phase: F_1.51,39.3_ = 27.3, p<0.0001; interaction: F_1.51,39.3_ = 0.39, p = 0.62, Fig. 3A.).

**Fig. 3.**
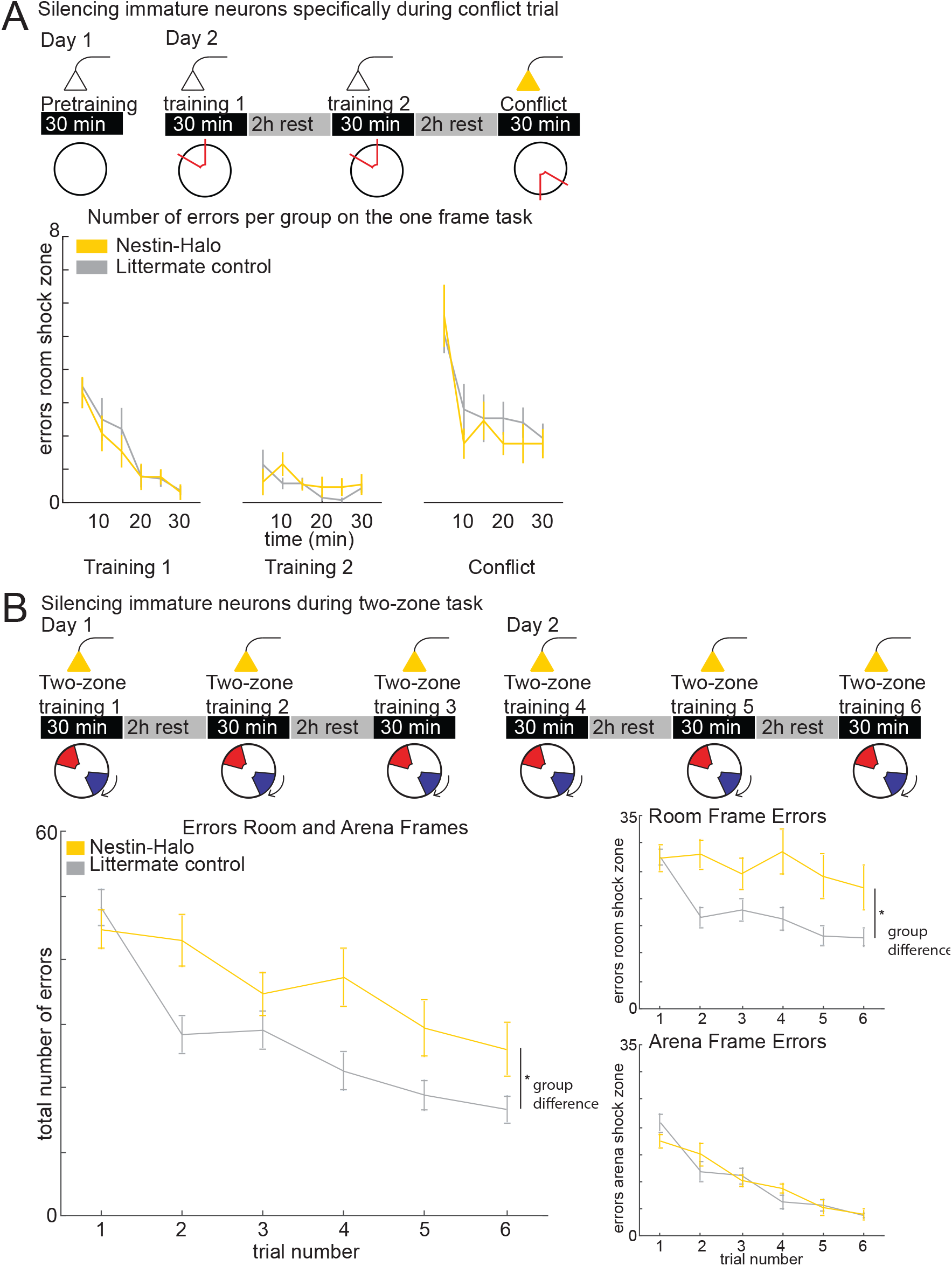
When mice did not have laser inhibition of immature neurons during initial learning of the stationary shock zone, mice with subsequently light-silenced immature granule cells are later impaired on a memory discrimination task with large cognitive load, specifically in avoiding the stationary shock zone. A) With the laser on only during the Conflict trial, NestinCre^+^ mice with optogenetcally-silenced immature cells are not impaired on the PIC protocol. Initial training 1: Genotype x Time repeated measures ANOVA; genotype: F_1,25_ = 0.28, p = 0.60; time: F_5,125_ = 25.4, p < 0.0001; interaction: F_5,8_ = 0.37, p = 0.87. Initial training 2: Genotype x Time repeated measures ANOVA; genotype: F_1,25_ = 0.26, p = 0.62; time: F_5,10_ = 3.51, p = 0.005; interaction: F_5,8_ = 1.91, p = 0.10. Conflict trial: Genotype x Time repeated measures ANOVA; genotype: F_1,25_ = 0.40, p = 0.53; time: F_5,125_ = 13.5, p < 0.0001; interaction: F_5,125_ = 0.63, p = 0.68. B) One week later, in the two-zone task with larger cognitive load, the same mice with silenced immature granule cells demonstrate decreased ability to discriminate between a previously learned stationary shock zone and a new qualitatively different rotating shock zone. Mice with light silenced immature neurons made more combined total number of stationary room frame and rotating arena frame errors (Genotype X Trial repeated measures ANOVA; genotype: F_1,26_ = 6.38, p=0.018; trial: F_5,22_ = 21.3, p < 0.0001; interaction: F5,22 = 3.73, p = 0.014), as well as more stationary room frame errors (Genotype X Trial repeated measures ANOVA; genotype: F_1,26_ = 8.43, p=0.007; trial: F_5,22_ = 5.72, p = 0.002; interaction: F_5,22_ = 5.12, p = 0.003). However, there were no genotype differences in the rotating arena frame errors (Genotype X Trial repeated measures ANOVA; genotype: F_1,26_ = 0.03, p=0.86; trial: F_5,22_ = 57.8, p < 0.0001; interaction: F_5,22_ = 1.17, p = 0.36).

Mice with X-ray or genetic ablation of immature neurons were impaired on the conflict trial in a similar PIC protocol (Burghardt et al., 2012), although in that study training was spaced over more days. We considered that laser light for silencing might not reach enough immature neurons, since the beam angle is approximately 60° and 100 μm above the dentate gyrus, thus light inhibiting an area of 100×100 μm. Accordingly, we hypothesized that more subtle behavioral differences between genotypes (with only a fraction of immature neurons compromised) could surface on a more demanding task.

To test if mice with light-silenced immature neurons were impaired on the two-zone task with larger cognitive load, NestinCre^+^-Halorhodopsin and control mice underwent an identical two-zone paradigm as the POMC mice, one week after training on the PIC protocol. During the two-zone task (Fig. 2B.), the groups of mice that had light-silenced neurons in all trials of the PIC paradigm, made similar amounts of stationary room frame shock zone errors. However, the NestinCre^+^-Halorhodopsin mice made more rotating arena frame shock zone errors compared to the NestinCre^−^ controls. Thus, the memory discrimination impairment after light-silencing immature granule cells is similar to the impairments in the POMC-Cre^+^-Halo mice.

Interestingly, for the cohort that was trained on the PIC protocol with the laser on only during the Conflict trial (Fig. 3), NestinCre^+^-Halorhodopsin mice with silenced immature neurons took longer to learn the two-zone task as indicated by more stationary room-frame zone avoidance errors up until the end of training, but the impairment was not observed in rotating arena-frame zone errors (Fig. 3B).

These data suggest that immature neurons are important for memory discrimination when the cognitive load is large and the memory distinction that has to be made is qualitative within the same trial (stationary versus rotating). Furthermore, the differences in which shock zone was better avoided (Fig. 2B vs. Fig. 3B) indicate that previous experience modulates the role that immature neurons may play.

The active place avoidance task relies heavily on information about space, like the signals that are carried by the medial entorhinal cortical (MEC) afferents. However, entorhinal cortex innervation of immature neurons comes predominantly from the lateral entorhinal cortex (Vivar et al., 2012; Vivar and van Praag, 2013; Woods et al., 2018), which preferentially targets the suprapyramidal blade of the DG (Witter, 2007). Distinct from MEC spatial signals, the discharge of LEC cells appears to signal non-spatial information such as context and objects (Deshmukh and Knierim, 2011; Knierim et al., 2014; Suzuki et al., 1997; Tsao et al., 2013) along with perirhinal cortical inputs that also preferentially target immature DG granule cells (Vivar et al., 2012; Vivar and van Praag, 2013). This functional anatomy, suggests that immature DG cells may be especially important for object-location associations and discriminations.

To investigate whether immature neurons (<5 weeks post-mitosis) contribute to the ability to discriminate objects, seven male NestinCreERT2^+^-Arch mice and seven male littermate NestinCreERT2^−^-Arch control mice performed a novel object discrimination task and an object displacement discrimination task. While control mice discriminated between the novel (Fig. 4A) and displaced objects (Fig. 4B), preferring the altered objects, the mice with laser-silenced immature neurons, did not discriminate between the objects or locations consistent with previous findings (Goodman et al., 2010; Suárez - Pereira et al., 2014), although when compared to each other, the genotypes did not differ on either the novel object task (t_12_ = 0.94, p = 0.37) nor the object displacement task (t_12_ = 1.0, p = 0.34). Importantly, these data also demonstrate that light-silencing immature neurons in the present conditions is sufficient to impair performance on memory discrimination tasks.

**Fig. 4.**
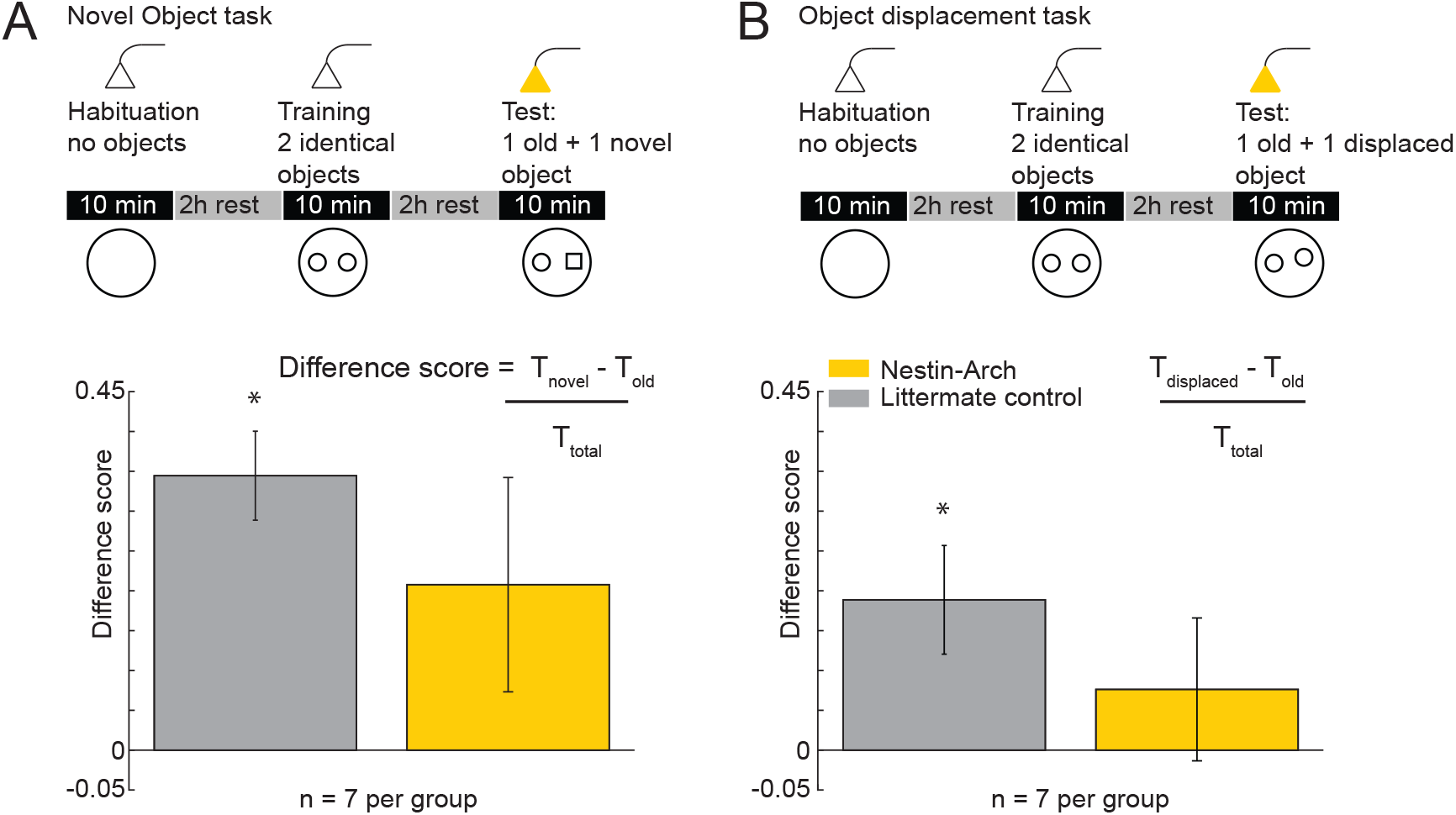
Optogenetic silencing of immature neurons impairs discriminating between novel and displaced objects. Unlike control mice, mice with silenced immature neurons do not discriminate between A) a familiar and a novel object (control mice: t_6_ = 6.17, p < 0.0001; Nestin-Cre^+^ mice: t_6_ =1.54, p = 0.17) or B) a familiarly-placed and a 4-cm displaced object (control mice: t_6_ = 2.77, p = 0.03; Nestin-Cre^+^ mice: (t_6_ = 0.85, p = 0.43).

## DISCUSSION

### Summary

Optogenetically silencing mouse DG granule cells impaired discrimination of memory for an initially-learned location of shock and a novel location of shock in the active place avoidance conflict task variant. Granule cell silencing also impaired performance of the two-zone task variant, further indicating that instead of causing a general spatial deficit, the manipulation of DG diminishes the ability to discriminate between memory of the previously learned stationary shock zone and the newly experienced rotating shock zone. In this case the DG manipulation impaired learning to avoid the new, rotating shock zone without impairing the familiar shock zone (Fig. 1). These findings provide additional support for the importance of the DG in performing memory discrimination in situations where the changes to the environment are subtle (Gilbert et al., 2001; Kheirbek et al., 2013; McHugh et al., 2007; Yassa and Stark, 2011).

In contrast to the effects of manipulating granule cells, selective inactivation of immature granule cells did not diminish memory discrimination tested with the conflict task variant, but the manipulation was sufficient to impair avoiding one of the two shock zones on the two-zone task variant. Importantly, which of the two simultaneous shock zones the experimental mice were deficient in avoiding, depended on whether or not immature neurons had been silenced during initial learning of the stationary shock zone. Because the deficit depends on whether or not immature granule cells were disturbed during initial learning, we take this as further evidence of a role in memory discrimination (Figs. 2,3). Inactivating immature neurons also caused deficits in discriminating novel and displaced objects, demonstrating a role of these cells in discriminating memories generally, not merely place memories (Fig. 4). The reversible lesion approach employed in this study replicates and extends previous findings of impaired performance in the two-zone task after permanent lesion of immature neurons (Burghardt et al., 2012), by the novel demonstration that consequences of silencing immature neurons during memory discrimination changes depending on an animal’s recent experience.

### Possible functional roles of immature DG granule cells

Immature neurons are highly excitable and plastic in comparison to mature granule cells, and they receive decreased inhibition and show more overlapping activity in response to perforant path stimulation and less discriminative spatial tuning *in vivo* (Danielson et al., 2016; Dieni et al., 2013; Kheirbek et al., 2012; Marin-Burgin et al., 2012). One “integrator” hypothesis for their role in memory processing is that immature neurons are more likely to (indiscriminately) encode new information and consequently link events that occur close in time (Aimone et al., 2010a; Deng et al., 2010). Because immature neurons are only highly excitable for a few weeks, they become less likely to be part of a new memory as they get older. Accordingly, immature neurons are optimized for encoding new information in a familiar context as well as separately encoding information about new contexts separated by time (Aimone et al., 2009; Rangel et al., 2014; Rangel et al., 2013). Indeed, immature neurons were specifically important for encoding in a novel context under circumstances of high interference (Danielson et al., 2016).

An alternative “inhibition” hypothesis is that immature neurons orchestrate inhibition onto mature neurons to sparsify memory representations. Several studies demonstrated increased excitability in the DG network after ablation or silencing of immature neurons, and decreased excitability of mature granule cells after optogenetic activation of immature neurons (Anacker et al., 2018; Burghardt et al., 2012; Drew et al., 2016; Ikrar et al., 2013; Lacefield et al., 2010; Park et al., 2015; Sahay et al., 2011b). The integrator and inhibition hypotheses are not mutually exclusive. Immature neurons could be directly involved in both the encoding and integration of temporally distinct information as well as the manipulation of mature granule cells for sparse and distinctive coding of information in memory (Anacker and Hen, 2017).

Our findings from spatially-restricted optogenetic inhibition of a subpopulation of DG neurons can be understood from both the integrator and inhibition hypotheses. Beginning with the inhibition hypothesis, if the role of immature neurons is indirect synaptic inhibition of mature granule cells, then silencing immature neurons is disinhibitory, resulting in increased mature granule cell excitability and consequently a less sparse network representation by DG granule cells that will tend to be excessively co-active. Such overexcitability might degrade the stored memories of the stationary frame locations of shock in the mice that learned these locations without laser inhibition during the PIC protocol, as observed (Fig. 3). Immature granule cell ablation results in activity-dependent dysregulation of inhibition (Park et al., 2015), and if this is effectively disinhibitory (Burghardt et al., 2012), then cell pairs that were initially weakly coactive are more likely to become aberrantly coactivate. Such aberrant coactivation has been shown in both modelling and experimental studies to degrade established place avoidance ensemble representations (Kao et al., 2017; Olypher et al., 2006). On the surface, it is unclear why new, rotating frame locations can be encoded in this overactive, disinhibited state, as we observed (Fig. 3), however such coactivity increases are demonstrated in network models to spare learning new representations, while impairing switching between representations (Olypher et al., 2006). Combined with a degraded representation of the stationary room locations of shock, these features would favor learning new rotating arena-frame locations during inhibition dysregulation, and arena-frame representations would dominate the corrupted room-frame representations. Given we only assessed behavior, these conjectures on neural activity are necessarily speculative and merit direct investigation in future work. Nonetheless, the pattern of behavioral results unambiguously indicates that the consequences for behavior depend on the history of learning and cell-specific inactivation. These complexities suggest and in fact favor dynamic neural network accounts, consistent with recent findings that demonstrate that DG place cell discharge network interactions, but not changes in single cell firing, mirror successful memory discrimination in the conflict task variant of the active place avoidance task (van Dijk and Fenton, 2018).

The present findings are also in line with an integrator role for immature DG granule cells. In this view, immature neurons are likely to be incorporated into the learned representations of the stationary locations of shock in the mice that had no laser inhibition during Initial training. During the subsequently trained two-zone task, a subpopulation of these immature neurons is silenced and retrieval of the stationary shock zone is disrupted as observed (Fig. 3), and as also seen after ablation of immature neurons post-encoding (Arruda-Carvalho et al., 2011; Gu et al., 2012).

Results from the two-zone tasks are similar between the POMC-Cre^+^ mice and the NestinCreERT+ mice, even though ten times fewer neurons should be silenced in the latter case of the immature neurons. This provides evidence for a preferential role of immature granule cells in encoding and discrimination of similar memories. An early version of the integrator hypothesis predicted that mature granule cells were not involved in encoding and discrimination of similar memories (Aimone et al., 2010b; Alme et al., 2010), but the current results do not support that hypothesis (Fig. 1).

In summary, these data as well as data from published work, together demonstrate roles for both mature and immature DG granule cells in memory discrimination. Although optogenetic silencing did not inhibit the entire population of targeted neurons, the findings are nonetheless, not easily explained by an exclusive role of immature granule cells in memory encoding or discrimination. Instead, the findings point to roles in potentially time-varying, dynamic network interactions amongst the various cell types in the DG that include mossy cells and a variety of inhibitory interneurons, as well as mature and immature granule cells.

## ACKNOWLEDGMENTS

Supported by NIH grant R01AG043688. We are grateful to Mazen Kheirbek for providing Nestin-CreERT2 mice and helpful discussions.

## REFERENCES

Aimone JB, Deng W, Gage FH. 2010a. Adult neurogenesis: integrating theories and separating functions. Trends Cogn Sci 14(7):325–37.

Aimone JB, Deng W, Gage FH. 2010b. Put them out to pasture? What are old granule cells good for, anyway…? Hippocampus 20(10):1124–5.

Aimone JB, Wiles J, Gage FH. 2009. Computational influence of adult neurogenesis on memory encoding. Neuron 61(2):187–202.

Alme CB, Buzzetti RA, Marrone DF, Leutgeb JK, Chawla MK, Schaner MJ, Bohanick JD, Khoboko T, Leutgeb S, Moser EI and others. 2010. Hippocampal granule cells opt for early retirement. Hippocampus 20(10):1109–23.

Amaral DG, Ishizuka N, Claiborne B. 1990. Neurons, numbers and the hippocampal network. Prog Brain Res 83:1–11.

Amaral DG, Scharfman HE, Lavenex P. 2007. The dentate gyrus: fundamental neuroanatomical organization (dentate gyrus for dummies). Prog Brain Res 163:3–22.

Anacker C, Hen R. 2017. Adult hippocampal neurogenesis and cognitive flexibility - linking memory and mood. Nat Rev Neurosci 18(6):335–346.

Anacker C, Luna VM, Stevens GS, Millette A, Shores R, Jimenez JC, Chen B, Hen R. 2018. Hippocampal neurogenesis confers stress resilience by inhibiting the ventral dentate gyrus. Nature.

Arruda-Carvalho M, Sakaguchi M, Akers KG, Josselyn SA, Frankland PW. 2011. Posttraining ablation of adult-generated neurons degrades previously acquired memories. J Neurosci 31(42):15113–27.

Bures J, Buresova O. 1990. Reversible lesions allow reinterpretation of system level studies of brain mechanisms of behavior. Concepts in Neuroscience 1:69–89.

Burghardt NS, Park EH, Hen R, Fenton AA. 2012. Adult-born hippocampal neurons promote cognitive flexibility in mice. Hippocampus 22(9):1795–808.

Cimadevilla JM, Fenton AA, Bures J. 2001. New spatial cognition tests for mice: passive place avoidance on stable and active place avoidance on rotating arenas. Brain Res Bull 54(5):559–63.

Clelland CD, Choi M, Romberg C, Clemenson GD, Jr., Fragniere A, Tyers P, Jessberger S, Saksida LM, Barker RA, Gage FH and others. 2009. A functional role for adult hippocampal neurogenesis in spatial pattern separation. Science 325(5937):210–3.

Coulter DA, Carlson GC. 2007. Functional regulation of the dentate gyrus by GABA-mediated inhibition. Prog Brain Res 163:235–43.

Danielson Nathan B, Kaifosh P, Zaremba Jeffrey D, Lovett-Barron M, Tsai J, Denny Christine A, Balough Elizabeth M, Goldberg Alexander R, Drew Liam J, Hen R and others. 2016. Distinct Contribution of Adult-Born Hippocampal Granule Cells to Context Encoding. Neuron 90(1):101–112.

Deng W, Aimone JB, Gage FH. 2010. New neurons and new memories: how does adult hippocampal neurogenesis affect learning and memory? Nat Rev Neurosci 11(5):339–50.

Deshmukh SS, Knierim JJ. 2011. Representation of non-spatial and spatial information in the lateral entorhinal cortex. Front Behav Neurosci 5:69.

Dieni CV, Nietz AK, Panichi R, Wadiche JI, Overstreet-Wadiche L. 2013. Distinct Determinants of Sparse Activation during Granule Cell Maturation. The Journal of Neuroscience 33(49):19131–19142.

Dranovsky A, Picchini AM, Moadel T, Sisti AC, Yamada A, Kimura S, Leonardo ED, Hen R. 2011. Experience dictates stem cell fate in the adult hippocampus. Neuron 70(5):908–23.

Drew LJ, Kheirbek MA, Luna VM, Denny CA, Cloidt MA, Wu MV, Jain S, Scharfman HE, Hen R. 2016. Activation of local inhibitory circuits in the dentate gyrus by adult-born neurons. Hippocampus 26(6):763–778.

Ge S, Yang CH, Hsu KS, Ming GL, Song H. 2007. A critical period for enhanced synaptic plasticity in newly generated neurons of the adult brain. Neuron 54(4):559–66.

Gilbert PE, Kesner RP, Lee I. 2001. Dissociating hippocampal subregions: double dissociation between dentate gyrus and CA1. Hippocampus 11(6):626–36.

Goodman T, Trouche S, Massou I, Verret L, Zerwas M, Roullet P, Rampon C. 2010. Young hippocampal neurons are critical for recent and remote spatial memory in adult mice. Neuroscience 171(3):769–778.

Gu Y, Arruda-Carvalho M, Wang J, Janoschka SR, Josselyn SA, Frankland PW, Ge S. 2012. Optical controlling reveals time-dependent roles for adult-born dentate granule cells. Nat Neurosci 15(12):1700–6.

Ikrar T, Guo N, He K, Besnard A, Levinson S, Hill A, Lee HK, Hen R, Xu X, Sahay A. 2013. Adult neurogenesis modifies excitability of the dentate gyrus. Front Neural Circuits 7:204.

Kao HY, Dvorak D, Park E, Kenney J, Kelemen E, Fenton AA. 2017. Phencyclidine discoordinates hippocampal network activity but not place fields. J Neurosci 37:12031–12049.

Kelemen E, Fenton AA. 2010. Dynamic grouping of hippocampal neural activity during cognitive control of two spatial frames. PLoS Biol 8(6):e1000403.

Kelemen E, Fenton AA. 2016. Coordinating different representations in the hippocampus. Neurobiol Learn Mem 129:50–9.

Kheirbek MA, Drew LJ, Burghardt NS, Costantini DO, Tannenholz L, Ahmari SE, Zeng H, Fenton AA, Hen R. 2013. Differential Control of Learning and Anxiety along the Dorsoventral Axis of the Dentate Gyrus. Neuron 77(5):955–68.

Kheirbek MA, Tannenholz L, Hen R. 2012. NR2B-dependent plasticity of adult-born granule cells is necessary for context discrimination. J Neurosci 32(25):8696–702.

Knierim JJ, Neunuebel JP, Deshmukh SS. 2014. Functional correlates of the lateral and medial entorhinal cortex: objects, path integration and local-global reference frames. Philos Trans R Soc Lond B Biol Sci 369(1635):20130369.

Lacefield CO, Itskov V, Reardon T, Hen R, Gordon JA. 2010. Effects of adult-generated granule cells on coordinated network activity in the dentate gyrus. Hippocampus 22(1):106–16.

Lee I, Hunsaker MR, Kesner RP. 2005. The role of hippocampal subregions in detecting spatial novelty. Behav Neurosci 119(1):145–53.

Marin-Burgin A, Mongiat LA, Pardi MB, Schinder AF. 2012. Unique processing during a period of high excitation/inhibition balance in adult-born neurons. Science 335(6073):1238–42.

Marr D. 1971. Simple memory: a theory for archicortex. Philos Trans R Soc Lond B Biol Sci 262(841):23–81.

McHugh TJ, Jones MW, Quinn JJ, Balthasar N, Coppari R, Elmquist JK, Lowell BB, Fanselow MS, Wilson MA, Tonegawa S. 2007. Dentate gyrus NMDA receptors mediate rapid pattern separation in the hippocampal network. Science 317(5834):94–9.

Mody I. 2005. Aspects of the homeostaic plasticity of GABAA receptor-mediated inhibition. J Physiol 562(Pt 1):37–46.

Mulders WH, West MJ, Slomianka L. 1997. Neuron numbers in the presubiculum, parasubiculum, and entorhinal area of the rat. J Comp Neurol 385(1):83–94.

Nakashiba T, Cushman JD, Pelkey KA, Renaudineau S, Buhl DL, McHugh TJ, Rodriguez Barrera V, Chittajallu R, Iwamoto KS, McBain CJ and others. 2012. Young dentate granule cells mediate pattern separation, whereas old granule cells facilitate pattern completion. Cell 149(1):188–201.

Olypher AV, Klement D, Fenton AA. 2006. Cognitive disorganization in hippocampus: a physiological model of the disorganization in psychosis. J Neurosci 26(1):158–68.

Overstreet-Wadiche LS, Westbrook GL. 2006. Functional maturation of adult-generated granule cells. Hippocampus 16(3):208–15.

Park EH, Burghardt NS, Dvorak D, Hen R, Fenton AA. 2015. Experience-Dependent Regulation of Dentate Gyrus Excitability by Adult-Born Granule Cells. J Neurosci 35(33):11656–66.

Rangel LM, Alexander AS, Aimone JB, Wiles J, Gage FH, Chiba AA, Quinn LK. 2014. Temporally selective contextual encoding in the dentate gyrus of the hippocampus. Nat Commun 5:3181.

Rangel LM, Quinn LK, Chiba AA, Gage FH, Aimone JB. 2013. A hypothesis for temporal coding of young and mature granule cells. Front Neurosci 7:75.

Sahay A, Scobie KN, Hill AS, O’Carroll CM, Kheirbek MA, Burghardt NS, Fenton AA, Dranovsky A, Hen R. 2011a. Increasing adult hippocampal neurogenesis is sufficient to improve pattern separation. Nature 472(7344):466–70.

Sahay A, Wilson DA, Hen R. 2011b. Pattern separation: a common function for new neurons in hippocampus and olfactory bulb. Neuron 70(4):582–8.

Schmidt-Hieber C, Jonas P, Bischofberger J. 2004. Enhanced synaptic plasticity in newly generated granule cells of the adult hippocampus. Nature 429(6988):184–7.

Senzai Y, Buzsáki G. 2017. Physiological Properties and Behavioral Correlates of Hippocampal Granule Cells and Mossy Cells. Neuron 93(3):691–704.e5.

Suárez - Pereira I, Canals S, Carrión Ángel M. 2014. Adult newborn neurons are involved in learning acquisition and long-term memory formation: The distinct demands on temporal neurogenesis of different cognitive tasks. Hippocampus 25(1):51–61.

Suzuki WA, Miller EK, Desimone R. 1997. Object and Place Memory in the Macaque Entorhinal Cortex. Journal of Neurophysiology 78(2):1062–1081.

Talbot ZN, Sparks FT, Dvorak D, Curran BM, Alarcon JM, Fenton AA. 2018. Normal CA1 Place Fields but Discoordinated Network Discharge in a Fmr1-Null Mouse Model of Fragile X Syndrome. Neuron.

Toni N, Laplagne DA, Zhao C, Lombardi G, Ribak CE, Gage FH, Schinder AF. 2008. Neurons born in the adult dentate gyrus form functional synapses with target cells. Nat Neurosci 11(8):901–7.

Treves A, Rolls ET. 1994. Computational analysis of the role of the hippocampus in memory. Hippocampus 4(3):374–91.

Tsao A, Moser MB, Moser EI. 2013. Traces of experience in the lateral entorhinal cortex. Curr Biol 23(5):399–405.

van Dijk MT, Fenton AA. 2018. On How the Dentate Gyrus Contributes to Memory Discrimination. Neuron 98(4):832–845.e5.

Vivar C, Potter MC, Choi J, Lee JY, Stringer TP, Callaway EM, Gage FH, Suh H, van Praag H. 2012. Monosynaptic inputs to new neurons in the dentate gyrus. Nat Commun 3:1107.

Vivar C, van Praag H. 2013. Functional circuits of new neurons in the dentate gyrus. Front Neural Circuits 7:15.

Witter MP. 2007. The perforant path: projections from the entorhinal cortex to the dentate gyrus. Prog Brain Res 163:43–61.

Woods NI, Vaaga CE, Chatzi C, Adelson JD, Collie MF, Perederiy JV, Tovar KR, Westbrook GL. 2018. Preferential Targeting of Lateral Entorhinal Inputs onto Newly Integrated Granule Cells. J Neurosci 38(26):5843–5853.

Yassa MA, Stark CE. 2011. Pattern separation in the hippocampus. Trends Neurosci 34(10):515–25.

Zhao C, Teng EM, Summers RG, Jr., Ming GL, Gage FH. 2006. Distinct morphological stages of dentate granule neuron maturation in the adult mouse hippocampus. J Neurosci 26(1):3–11.

